# Identification of inflammatory biomarkers in IgA nephropathy using the NanoString technology: a validation study

**DOI:** 10.1101/2023.09.11.556864

**Authors:** Laurence Gaumond, Caroline Lamarche, Stéphanie Beauchemin, Nathalie Henley, Naoual Elftouh, Casimiro Gerarduzzi, Louis-Philippe Laurin

## Abstract

**Objective and design:** Immunoglobulin A nephropathy (IgAN) is a kidney disease characterized by the accumulation of IgA deposits in the glomeruli of the kidney, leading to inflammation and damage to the kidney. The inflammatory markers involved in IgAN remain to be defined. Gene expression analysis platforms, such as the NanoString nCounter system, are promising screening and diagnostic tools, especially in the oncology field, but its role as a diagnostic and prognostic tool in IgAN remains scarce. In this study, we aimed to validate the use of NanoString technology to identify potential inflammatory biomarkers involved in the progression of IgAN.

**Subjects:** A total of 30 patients with biopsy-proven IgAN and 7 cases of antineutrophil cytoplasmic antoantibodies (ANCA) associated pauci-immune glomerulonephritis were included for gene expression measurement. For the immunofluorescence validation experiments, a total of 6 IgAN patients and 3 healthy controls were included.

**Methods:** Total RNA was extracted from formalin-fixed-paraffin-embedded kidney biopsy specimens, and a customized 48-plex human gene CodeSet was used to study 29 genes implicated in different biological pathways. Comparisons in gene expression were made between IgAN and ANCA-associated pauci-immune glomerulonephritis patients to delineate an expression profile specific to IgAN. Gene expression was compared between patients with low and moderate risk of progression. Genes for which RNA expression was associated with disease progression were analyzed for protein expression by immunofluorescence, and compared with healthy controls.

**Results:** IgAN patients had a distinct gene expression profile with decreased expression in genes *IL-6, INFG* and *C1QB* compared to ANCA patients. *C3* and *TNFR2* were identified as potential biomarkers for IgAN progression in patients early in their disease course. Protein expression for those 2 candidate genes was upregulated in IgAN patients compared to controls. Expression of genes implicated in fibrosis (*PTEN, CASPASE 3, TGM2, TGFB1, IL2*, and *TNFRSF1B*) was more pronounced in IgAN patient with severe fibrosis compared to those with none.

**Conclusions:** Our findings validate our NanoString mRNA profiling by examining protein expression levels of two candidate genes, *C3* and *TNFR2*, in IgAN patients and healthy controls. We were also able to identify several upregulated mRNA transcripts implicated in the development of fibrosis that may be considered as fibrotic markers within IgAN patients.

## INTRODUCTION

Immunoglobin A nephropathy (IgAN) is a complicated autoimmune condition where immunoglobulin A (IgA) is deposited in the kidney’s glomeruli, causing inflammation, and ultimately leading to kidney failure [1]. IgAN is a leading cause of primary glomerulonephritis and chronic kidney disease [2]. Several clinical factors have been identified over the last decades to help predict disease progression, such as sustained proteinuria over a gram a day, baseline estimated glomerular filtration rate (eGFR), high blood pressure and histopathologic findings from the Oxford classification [3]. In IgAN, the search for inflammatory biomarkers that can be utilized to track therapy effectiveness and foretell disease development remains unfinished [4]. No one biomarker can give a comprehensive picture of IgAN, so the use of several biomarkers may be required to correctly diagnose and treat the disease. Although a number of clinical indicators have been found to help predict disease progression, specific biomarkers involved in the underlying inflammatory processes are not integrated into any prediction tool. Therefore, further research is needed to validate the clinical utility of inflammatory biomarker panel in IgAN.

State-of-the-art platforms for gene expression analysis, such as the NanoString nCounter system, are promising screening and diagnostic tools, especially in the oncology field. This innovative process, which directly measures individual micro ribonucleic acid (RNA) transcripts without enzymatic steps, stands out from previous methods by its high sensitivity, precision, and efficiency [5]. Specifically, the NanoString nCounter system uses a digital molecular barcoding technology, which allows for high-throughput and sensitive detection of specific RNA molecules in a sample without amplification. It is often used in research to study gene expression, biomarker discovery, and pathway analysis. Increasingly used in several clinical settings, this technology could help identify biomarkers playing a role in various inflammatory pathologic processes.

This pilot study successfully validates the application of NanoString technology in identifying potential inflammatory targets related to different stages of IgAN disease progression. Our results revealed a distinctive gene expression profile in IgAN patients, notably, we identified *C3* and *TNFR2* as promising biomarkers for monitoring IgAN progression, with their protein expression being significantly upregulated in IgAN patients when compared to healthy controls. Additionally, our investigation uncovered heightened expression of genes associated with fibrosis in IgAN patients with severe fibrosis compared to those without fibrosis. This finding suggests the potential utility of these mRNA transcripts as fibrotic markers for IgAN patients. The successful validation of NanoString mRNA profiling and the identification of relevant inflammatory and fibrotic markers may pave the way for improved diagnostic and prognostic tools in IgAN, enhancing our understanding of the disease’s underlying inflammatory processes.

## METHODS

### Study design and population

All patients with biopsy-proven IgAN and antineutrophil cytoplasmic antoantibodies (ANCA) associated pauci-immune glomerulonephritis between 2005 and 2021 diagnosed at Maisonneuve-Rosemont Hospital, a single tertiary care hospital in Montreal, were considered for this retrospective study. Only patients with formalin-fixed-paraffin-embedded (FFPE) biopsy samples available for gene expression analysis were included. Patients with known secondary causes of IgAN were excluded. For the immunofluorescence experiments, healthy controls with no glomerular disease were recruited among patients who underwent a nephrectomy in the context of a kidney lesion. All subjects provided written informed consent. The study was approved by our institution’s Research Ethics Committee (N° MP-12-2014-525), in agreement with the Declaration of Helsinki.

### Data collection

Clinical data were collected by reviewing medical records, and included demographics, kidney presentation, comorbidities, history of familial kidney disease, and medications. Specifically, serum creatinine, eGFR and proteinuria were noted at the time of the biopsy, at 6 months from biopsy time and then annually to the last available follow-up visit and/or initiation renal replacement therapy. The Oxford MEST-C score [6] was retrieved from the biopsy report. Baseline eGFR was estimated using the Chronic Kidney Disease - Epidemiology Collaboration equation [7]. Urinary protein excretion was estimated using either total 24-hour proteinuria (in g/day), spot urinary protein-creatinine ratio (in mg/mmol) or spot urinary albumin-creatinine ratio (in mg/mmol). For IgAN, the risk of a 50% decline in eGFR or progression to end-stage kidney disease (ESKD) 5 years after kidney biopsy was calculated using the New International Risk-Prediction Tool in IgAN [8], a prediction model that includes baseline clinical and pathological characteristics, demographics, and medication use. The studied hard outcomes were ESKD (eGFR <15 ml/min/1.73m^2^, dialysis or kidney transplantation) or death.

### Gene expression measurement

For our NanoString analysis, a total 30 biopsy-confirmed IgA and 7 ANCA-positive pauci-immune glomerulonephritis patients were used. Total RNA from FFPE kidney biopsy specimens from patients with IgAN and ANCA-positive pauci-immune glomerulonephritis (control group) were extracted and purified using RNeasy FFPE kit (Qiagen) according to the manufacturer’s protocol. A total of 100 nanograms of RNA from each sample was hybridized to a customized 48-plex human gene CodeSet and processed on the nCounter platform. Purified RNA samples were analyzed using the nCounter gene expression system from NanoString Technology. A total of 29 genes implicated in different biological pathways were studied; those genes have been selected for their role in complement activation (*C2, C3, C1QA, C1QB*), apoptosis (*BAX, BAD, Caspase 3*), cell activation and/or leucocytes stimulation (*AGTR1, AGTR2, CXCL1, CD89S, ICAM1, MIF, PTEN, CD71, TGM2, TNFR1B, TNFR2, TNFSF13, IL-2, IL-4, IL-5, IL-6, IL-8, IL-10, IL-13, IFN-γ, TGF-β1, TNF*). Housekeeping gene (*ACTB, GAPDH, HPRT1, LDHA*) normalization was performed to adjust counts of all probes relative to a probe that are not expected to vary between samples; factor should be in the range between 0.1 and 10. All normalization procedures were performed according to the manufacturer’s protocol.

Data collection for gene expression was carried out in the nCounter Digital Analyzer. At the highest standard resolution, 555-1155 field of view were collected per flow cell using a microscope objective and a CCD camera yielding data of hundreds of thousands of target molecule counts. Every RNA target is identified by the color code generated by the ordered fluorescent segments present on the reporter probe. The expression level of each gene is determined by scoring the number of times its corresponding color code is detected. Data was imported into nSolver Analysis Software for downstream analysis.

Comparisons in gene expression were made between IgAN patients and those with another inflammatory glomerulonephritis (ANCA-associated pauci-immune glomerulonephritis) to depict an expression profile specific to IgAN. Subsequently, among individuals with IgAN, gene expression was compared between patients with low chance of progression to ESKD (<10%) and those at mild to moderate risk of progression (10-20%) with relatively minimal fibrosis on kidney biopsy. Genes for which RNA expression was associated with disease progression were analyzed in 6 patients for protein expression by immunofluorescence, and compared with healthy controls.

### Histology

Histologic slides for all patients with IgAN were reviewed by the same nephropathologist to provide MEST-C score: mesangial hypercellularity (with M0 and M1 corresponding to ≤50% and >50% of glomeruli with hypercellularity, respectively), segmental glomerulosclerosis (with S0 if absent and S1 if present), endocapillary hypercellularity (with E0 if absent and E1 if present), tubular atrophy/interstitial fibrosis (with T0, T1 and T2 corresponding to ≤25%, 26%-50% and >50% of cortical area involvement, respectively) and cellular/fibro cellular crescents (with C0, C1 and C2 corresponding to their absence, presence in ≥1 and <25% of glomeruli, and presence in ≥25% of the glomeruli, respectively). ANCA-associated pauci-immune glomerulonephritis was also graded based on its degree of tubular atrophy/interstitial fibrosis.

### Immunofluorescence

The kidneys were fixed in 10% formaldehyde, dehydrated, and embedded in paraffin. The tissue was sectioned (5 μM) and was subjected to antigen retrieval in citrate solution at pH 6. The sections were blocked with anti-donkey serum 5% and labelled with anti-Complement C3 (Catalog #PA1-29715 from Life Technologies/ThermoFisher) and TNFRSF1B (Catalog #MA5-31661 from Life Technologies/ThermoFisher). The slides were subsequently exposed to donkey anti-rabbit AF647-conjugated (1:400, Catalog # 711 605 152 Jackson ImmunoResearch Laboratories), donkey anti-mouse AF568 (Catalog # A10037 Life Technologies/ThermoFisher) and donkey anti-goat AF488 (Catalog # A11055 Life Technologies/ThermoFisher). Fluoroshield with DAPI (Millipore-Sigma) was used for nuclear staining and mounting. Slides were imaged using a Zeiss AxioObserver.Z1 inverted microscope coupled to an X-Cite 120LED Boost High-Power LED illumination system. Images for quantitative analysis were captured with a 20X objective and the number of positive cells was determined as the average of positive cells in at least 8 fields per kidney section.

### Statistical analysis

Baseline characteristics, clinical and pathological parameters, and gene expression were compared between patients with IgAN and those with ANCA-associated pauci-immune glomerulonephritis (controls). Among patients with IgAN, gene expression was compared between progression score groups (<10%, 10-20%, 21-40% and >40%), and more specifically between patients with score <10% and those with 10-20% to delineate markers of early disease. Wilcoxon or *t*-tests for continuous variables and the χ^2^ test or the Fisher exact tests were used for categorical variables. The Kruskal Wallis test was also used for comparison between more than two groups. Differences were considered statistically significant if p-value <0.05. Statistical analysis was performed by N.E. using SAS 9.4 (SAS Institute, Cary, NC).

## RESULTS

### Patient characteristics

Patients with IgAN with gene expression analyzed by NanoString were mostly Caucasians (76.7%) and females (73.3%), as shown in Table 1. Mean age at biopsy was 44.7±15.3 years. Mean baseline eGFR and proteinuria were 60.9±35.8 ml/min/1.73m^2^, and 3.2±2.9 g/d respectively; most patients (96.5%) had a stage 0 or 1 of fibrosis at diagnosis. At baseline, 60% of patients had chronic kidney disease stage 3 or 4. Treatments were highly variable, but a great proportion (80%) were on renin angiotensin aldosterone (RAA) system inhibitors and 45% received glucocorticoids at some point in their disease course. Patients with ANCA-associated pauci-immune glomerulonephritis were all Caucasians (100%), with a majority of females (57.1%). Mean age at biopsy was 62.6±9.4 years. At baseline, mean eGFR and proteinuria were 36.6±33 ml/min/1.73m^2^and 2.4±2.2 g/d respectively; all patients had stage 0 or 1 of fibrosis at diagnosis. Patients with ANCA-associated pauci-immune glomerulonephritis were significantly older and had more kidney impairment at baseline than patients with IgAN.

**Table 1.** Patient characteristics.

| Variable | ANCA-associated<br>pauci-immune<br>glomerulonephritis<br>(n=7) | IgA<br>nephropathy<br>(n=30) | P-value | N missing |
| --- | --- | --- | --- | --- |
| Female | 4 (57.1) | 22 (73.3) | 0.1827 |  |
| Age at biopsy, yrs | 62.6 (9.4) | 44.7 (15.3) | 0.0096 |  |
| BMI at baseline, kg/m<sup>2</sup> | 27.2 (4.4) | 28.3 (7.0) | 0.8442 | 4 |
| Ethnicity |  |  |  |  |
| Caucasian | 7 (100) | 23 (76.7) | 0.5694 |  |
| Afro-american | 0 | 1 (3.3) |  |  |
| Asian | 0 | 4 (13.3) |  |  |
| Others | 0 | 2 (6.7) |  |  |
| EGFR at baseline, ml/min per 1.73 m<sup>2</sup> | 36.6 (33.0) | 60.9 (35.8) | 0.0363 |  |
| Proteinuria at baseline, g/d | 2.4 (2.2) | 3.2 (2.9) | 0.4044 |  |
| Fibrosis on biopsy, % |  |  |  |  |
| T0 | 3 (50) | 11 (37.9) | 0.7975 | 2 |
| T1 | 3 (50) | 17 (58.6) |  |  |
| T2 | 0 | 1 (3.5) |  |  |
| Diabetes | 1 (14.3) | 9 (30) | 0.6471 |  |
| Hypertension | 6 (85.7) | 23 (76.7) | 1.0000 |  |
| Family history of ESKD | 0 | 1 (3.3) | 1.0000 |  |
| Smoking | 4 (57.1) | 15 (50) | 1.0000 |  |
| Medications* |  |  |  |  |
| RAAS inhibitors | 3 (42.9) | 24 (80) | 0.0688 |  |
| Glucocorticoids | 5 (83.3) | 9 (45) | 0.1696 | 11 |
| MMF | 1 (16.7) | 1 (5) | 0.4154 | 11 |
| Ballardie** regimen | 4 (66.7) | 1 (5) | 0.0047 | 11 |
| Cyclophosphamide | 3 (50.0) | 1 (5) | 0.0278 | 11 |
| Azathioprine | 2 (33.3) | 1 (5) | 0.1231 | 11 |
| Rituximab | 2 (33.3) | 0 (0) | 0.0462 | 11 |

Data are expressed as mean (standard deviation) or n (percent). ANCA, antineutrophil cytoplasmic autoantibody; IgA, immunoglobulin A; BMI, body mass index; EGFR, estimated glomerular filtration rate; ESKD, end-stage kidney disease; RAAS, renin-angiotensin-aldosterone system; MMF, mycophenolate mofetil
\*Received at any time during disease course.
\*\*Glucocorticoids and cyclophosphamide for the initial 3 months, then azathioprine for a minimum of 2 years

The 5-year risk of 50% decline in eGFR or ESKD after biopsy was variable in IgAN patients: 9, 5, 8 and 6 patients (2 missing) had a 5-year risk of progression of <10%, 10%-20%, 21%-40% and >40%, respectively. The median follow-up time for IgAN group was 4.62 years (interquartile range 2.45 years). Among patients with IgAN, 8 patients (26.7%) reached ESKD and 3 (10%) died.

### Gene expression profiles

IgAN patients had a different mRNA expression profile than patients with ANCA-associated pauci-immune glomerulonephritis. *IL-6, INFG* and *C1QB* were significantly transcribed at higher levels in patients with ANCA-associated pauci-immune glomerulonephritis (Table 2). *AGTR1* was more expressed in patients with IgAN compared to ANCA controls (p=0.05).

**Table 2.** Distribution of gene expression between IgAN and ANCA patients.

| Variable | ANCA |  | IgAN |  | P-value |
| --- | --- | --- | --- | --- | --- |
|  | n=7 | SD | n=30 | SD |  |
| *AGTR1* | 64.7 | 28.7 | 162.2 | 119.8 | 0.0470 |
| *AGTR2* | 3.2 | 2.0 | 9.3 | 12.6 | 0.1433 |
| *C1QA* | 42.3 | 36.5 | 19.8 | 13.8 | 0.0795 |
| *C1QB* | 1091.0 | 919.7 | 395.9 | 305.3 | 0.0180 |
| *C3* | 402.2 | 554.7 | 177.9 | 162.5 | 0.3356 |
| *CD71* | 103.8 | 45.5 | 154.5 | 103.7 | 0.3787 |
| *CD89S* | 3.0 | 1.9 | 3.3 | 2.2 | 0.9144 |
| *CXCL1* | 175.3 | 141.4 | 99.8 | 73.0 | 0.3355 |
| *ICAM1* | 197.7 | 147.5 | 127.8 | 83.1 | 0.2865 |
| *IFNG* | 6.5 | 7.7 | 1.9 | 2.0 | 0.0175 |
| *IL10* | 3.2 | 2.6 | 2.4 | 1.7 | 0.5118 |
| *IL13* | 4.8 | 2.6 | 3.5 | 2.6 | 0.2276 |
| *IL2* | 1.8 | 1.0 | 1.4 | 0.7 | 0.2471 |
| *IL4* | 1.7 | 1.2 | 1.1 | 0.4 | 0.0703 |
| *IL5* | 4.0 | 2.8 | 2.7 | 1.5 | 0.3690 |
| *IL6* | 9.7 | 5.3 | 3.4 | 3.1 | 0.0076 |
| *IL8* | 69.3 | 82.9 | 26.8 | 22.3 | 0.1896 |
| *MIF* | 1268.3 | 599.5 | 1384.8 | 631.1 | 0.7842 |
| *PTEN* | 390.3 | 237.1 | 470.0 | 241.6 | 0.5149 |
| *TGFB1* | 337.8 | 210.9 | 302.1 | 130.8 | 0.8168 |
| *TGM2* | 373.5 | 232.3 | 259.4 | 143.4 | 0.2345 |
| *TNF* | 41.7 | 37.4 | 27.8 | 19.3 | 0.5701 |
| *TNFR1B* | 434.0 | 231.9 | 447.7 | 251.1 | 0.8496 |
| *TNFRSF1B* | 140.2 | 96.1 | 102.9 | 50.8 | 0.2682 |
| *TNFSF13* | 82.5 | 42.8 | 75.1 | 29.3 | 0.5699 |
| *BAD* | 101.5 | 33.4 | 111.4 | 42.0 | 0.5013 |
| *BAX* | 321.0 | 168.4 | 311.0 | 160.5 | 0.8496 |
| *C2* | 125.2 | 115.3 | 73.1 | 68.4 | 0.3153 |
| *CASPASE3* | 111.2 | 83.3 | 93.2 | 54.7 | 0.8331 |
| *ACTB* | 7481.0 | 4373.3 | 7052.5 | 3307.5 | 0.8496 |
| *GAPDH* | 4209.8 | 1800.4 | 4779.9 | 3293.8 | 1.0000 |
| *HPRT1* | 235.5 | 117.2 | 238.7 | 135.1 | 0.8496 |
| *LDHA* | 1352.8 | 585.5 | 1346.0 | 815.7 | 0.9496 |

ANCA, antineutrophil cytoplasmic antoantibody; IgAN, immunoglobulin A nephropathy

Biomarkers distribution in function of the IgA nephropathy prediction tool score range were analyzed. Markers known as *CIQA, ICAM1, TNF* and *TNFSF1B* did not show a statistically significant distribution, but their distribution difference was near the significance level (Table 3). As shown in Table 4, *CIQB and TNFSF1B* (coding for TNFR2) expression were higher among patients with an IgA prediction tool score range of 10-20% compared to a score <10% (p-value = 0.01342 and p=0.03337, respectively). *C3* expression was also higher in patients with a score range of 10-20% (vs. with a score <10%) with a p-value of 0.04911.

**Table 3.** Distribution of gene expression by Risk-Prediction Tool score ranges.

| Variable | Score <10% |  | Score 10-20% |  | Score 20-40% |  | Score 40% |  | P-value* |
| --- | --- | --- | --- | --- | --- | --- | --- | --- | --- |
|  | n=9 | % | n=5 | % | n=8 | % | n=6 | % |  |
| *AGTR1* | 154.56 | 120.2 | 251.6 | 133.5 | 144.38 | 97.3 | 141.83 | 148.1 | 0.3845 |
| *AGTR2* | 6.78 | 6.4 | 4.4 | 4.2 | 13.88 | 21.9 | 10.67 | 9.7 | 0.7307 |
| *C1QA* | 14.11 | 8.9 | 36.4 | 17.0 | 18.5 | 10.7 | 17 | 13.8 | 0.0654 |
| *C1QB* | 255.33 | 156.5 | 736.8 | 565.1 | 390.63 | 148.2 | 355.5 | 212.2 | 0.1057 |
| *C3* | 116.11 | 124.2 | 318.4 | 205.0 | 159.63 | 118.2 | 150 | 143.2 | 0.1589 |
| *CD71* | 167.56 | 109.8 | 212 | 132.1 | 145.75 | 91.3 | 110.67 | 97.9 | 0.4605 |
| *CD89S* | 3.56 | 2.8 | 2.8 | 1.1 | 3.25 | 2.6 | 3.5 | 2.1 | 0.9565 |
| *CXCL1* | 73 | 68.0 | 113.2 | 47.8 | 111 | 78.9 | 116 | 96.8 | 0.6398 |
| *ICAM1* | 99 | 66.6 | 196.6 | 34.9 | 137.63 | 105.7 | 98.17 | 85.1 | 0.0838 |
| *IFNG* | 1.33 | 0.7 | 3.4 | 4.3 | 2 | 1.6 | 1.5 | 0.8 | 0.7648 |
| *IL10* | 2 | 1.1 | 2.6 | 1.1 | 3 | 2.0 | 1.33 | 0.5 | 0.1614 |
| *IL13* | 3.67 | 2.5 | 6 | 3.5 | 2.88 | 2.2 | 2.17 | 1.2 | 0.157 |
| *IL2* | 1.11 | 0.3 | 1.6 | 0.6 | 1.5 | 0.9 | 1.33 | 0.8 | 0.3551 |
| *IL4* | 1.22 | 0.7 | 1.2 | 0.5 | 1 | 0.0 | 1 | 0.0 | 0.5018 |
| *IL5* | 3 | 1.1 | 3.2 | 2.5 | 2.63 | 1.7 | 2.17 | 1.2 | 0.6339 |
| *IL6* | 3.67 | 3.3 | 5 | 3.2 | 2.13 | 1.4 | 2.33 | 2.0 | 0.2006 |
| *IL8* | 22.78 | 22.5 | 29.8 | 15.5 | 35.88 | 30.5 | 17.33 | 12.7 | 0.4965 |
| *MIF* | 1409.22 | 674.8 | 1796 | 439.1 | 1466.88 | 654.6 | 958.67 | 590.1 | 0.2334 |
| *PTEN* | 413.11 | 216.8 | 620.2 | 244.4 | 493.5 | 208.4 | 408.83 | 315.3 | 0.487 |
| *TGFB1* | 218.33 | 109.1 | 371.4 | 70.8 | 346.63 | 118.5 | 304.83 | 173.9 | 0.1202 |
| *TGM2* | 196.22 | 149.9 | 318 | 63.3 | 328.38 | 128.8 | 209.5 | 158.1 | 0.2137 |
| *TNF* | 17.33 | 10.7 | 40.8 | 16.2 | 32.88 | 19.6 | 26.83 | 27.7 | 0.0873 |
| *TNFR1B* | 394.33 | 201.7 | 682.2 | 285.1 | 440.25 | 220.9 | 334.17 | 252.8 | 0.2078 |
| *TNFRSF1B* | 70.78 | 36.8 | 140.8 | 47.0 | 116.63 | 36.0 | 99.67 | 67.5 | 0.0602 |
| *TNFSF13* | 62.33 | 31.0 | 82.4 | 8.2 | 90.75 | 30.0 | 66 | 35.0 | 0.2166 |
| *BAD* | 106.11 | 49.4 | 121.6 | 21.7 | 120 | 42.7 | 103.67 | 48.9 | 0.8908 |
| *BAX* | 274.78 | 137.4 | 424.8 | 135.6 | 319.63 | 155.6 | 251 | 181.1 | 0.2558 |
| *C2* | 79.11 | 88.4 | 91.4 | 68.0 | 76.88 | 70.3 | 41.5 | 25.0 | 0.5412 |
| *CASPASE3* | 72.56 | 39.3 | 134.4 | 58.9 | 102.38 | 46.3 | 78.67 | 70.8 | 0.1902 |

\*Kruskal-Wallis test for comparison between more than two groups. Two patients have the prediction tools missing.

**Table 4.** Mean difference of gene expression between Risk-Prediction Tool score groups <10% and 10-20%.

| Variable | Score <10% |  | Score 10-20% |  | Score <10 vs 10-20% |  | P-value |
| --- | --- | --- | --- | --- | --- | --- | --- |
|  | n=9 | SD | n=5 | SD | Mean diff | SE |  |
| *AGTR1* | 154.56 | 120.2 | 251.6 | 133.5 | 97.04 | 68.6 | 0.38664 |
| *AGTR2* | 6.78 | 6.4 | 4.4 | 4.2 | -2.38 | 7.4 | 0.97968 |
| *C1QB* | 255.33 | 156.5 | 736.8 | 565.1 | 481.47 | 155.0 | 0.01342 |
| *C3* | 116.11 | 124.2 | 318.4 | 205.0 | 202.29 | 79.9 | 0.04911 |
| *CD71* | 167.56 | 109.8 | 212 | 132.1 | 44.44 | 59.5 | 0.81152 |
| *CD89S* | 3.56 | 2.8 | 2.8 | 1.1 | -0.76 | 1.3 | 0.90161 |
| *CXCL1* | 73 | 68.0 | 113.2 | 47.8 | 40.20 | 42.1 | 0.67982 |
| *ICAM1* | 99 | 66.6 | 196.6 | 34.9 | 97.60 | 44.8 | 0.10232 |
| *IFNG* | 1.33 | 0.7 | 3.4 | 4.3 | 2.07 | 1.1 | 0.20489 |
| *IL10* | 2 | 1.1 | 2.6 | 1.1 | 0.60 | 0.8 | 0.78642 |
| *IL13* | 3.67 | 2.5 | 6 | 3.5 | 2.33 | 1.4 | 0.23824 |
| *IL2* | 1.11 | 0.3 | 1.6 | 0.6 | 0.49 | 0.4 | 0.47247 |
| *IL4* | 1.22 | 0.7 | 1.2 | 0.5 | -0.02 | 0.2 | 0.99947 |
| *IL5* | 3 | 1.1 | 3.2 | 2.5 | 0.20 | 0.9 | 0.99288 |
| *IL6* | 3.67 | 3.3 | 5 | 3.2 | 1.33 | 1.4 | 0.69613 |
| *IL8* | 22.78 | 22.5 | 29.8 | 15.5 | 7.02 | 12.7 | 0.90954 |
| *MIF* | 1409.22 | 674.8 | 1796 | 439.1 | 386.78 | 344.5 | 0.5679 |
| *PTEN* | 413.11 | 216.8 | 620.2 | 244.4 | 207.09 | 135.5 | 0.32601 |
| *TGFB1* | 218.33 | 109.1 | 371.4 | 70.8 | 153.07 | 68.8 | 0.09321 |
| *TGM2* | 196.22 | 149.9 | 318 | 63.3 | 121.78 | 75.2 | 0.28267 |
| *TNF* | 17.33 | 10.7 | 40.8 | 16.2 | 23.47 | 10.5 | 0.09101 |
| *TNFR1B* | 394.33 | 201.7 | 682.2 | 285.1 | 287.87 | 130.4 | 0.09666 |
| *TNFRSF1B* | 70.78 | 36.8 | 140.8 | 47.0 | 70.02 | 25.8 | 0.03337 |
| *TNFSF13* | 62.33 | 31.0 | 82.4 | 8.2 | 20.07 | 16.3 | 0.495 |
| *BAD* | 106.11 | 49.4 | 121.6 | 21.7 | 15.49 | 24.5 | 0.87348 |
| *BAX* | 274.78 | 137.4 | 424.8 | 135.6 | 150.02 | 85.0 | 0.22153 |
| *C2* | 79.11 | 88.4 | 91.4 | 68.0 | 12.29 | 39.2 | 0.98119 |
| *CASPASE3* | 72.56 | 39.3 | 134.4 | 58.9 | 61.84 | 29.3 | 0.11736 |
| *ACTB* | 5964.11 | 2891.5 | 9102.2 | 1809.4 | 3138.09 | 1843.7 | 0.24625 |
| *GAPDH* | 5045.22 | 2969.0 | 7142.6 | 4504.5 | 2097.38 | 1797.5 | 0.53886 |
| *HPRT1* | 236.78 | 126.2 | 311.8 | 112.5 | 75.02 | 77.3 | 0.66952 |
| *LDHA* | 1259.22 | 714.4 | 1744.2 | 626.0 | 484.98 | 470.8 | 0.62995 |

### Immunofluorescence

To ensure the accuracy and reliability of our findings obtained through Nanostring mRNA profiling, we devised a validation strategy that involved the examination of protein expression levels of candidate genes. We randomly selected two genes, namely Complement C3 and TNFR2, to conduct our protein expression analyses. To this end, we obtained biopsies from patients diagnosed with IgA nephropathy that had high degree of fibrosis compared to the healthy controls (Fig. 1A). Our findings revealed that both target genes identified in gene expression analysis (*C3* and *TNFSF1B*) exhibited a significant upregulation of protein expression in the biopsies obtained from patients with IgA nephropathy when compared to healthy tissue (Fig. 1B). Importantly, the expression patterns of Complement C3 and TNFR2 were distinct, further underscoring the significance of our observations. Complement C3 was found to be distributed around the tubular epithelial cells and glomerulus, with a modest signal, suggesting a possible role in the regulation of inflammation and immune responses is such regions. On the other hand, the signal for TNFR2 was more pronounced and concentrated in distinct regions that differed from those observed for Complement C3, highlighting its potential role in cellular proliferation and differentiation. Overall, our findings validate our NanoString mRNA profiling by examining protein expression levels of two candidate genes, *C3* and *TNFR2*, in IgAN patients and healthy controls.

**Figure 1.**
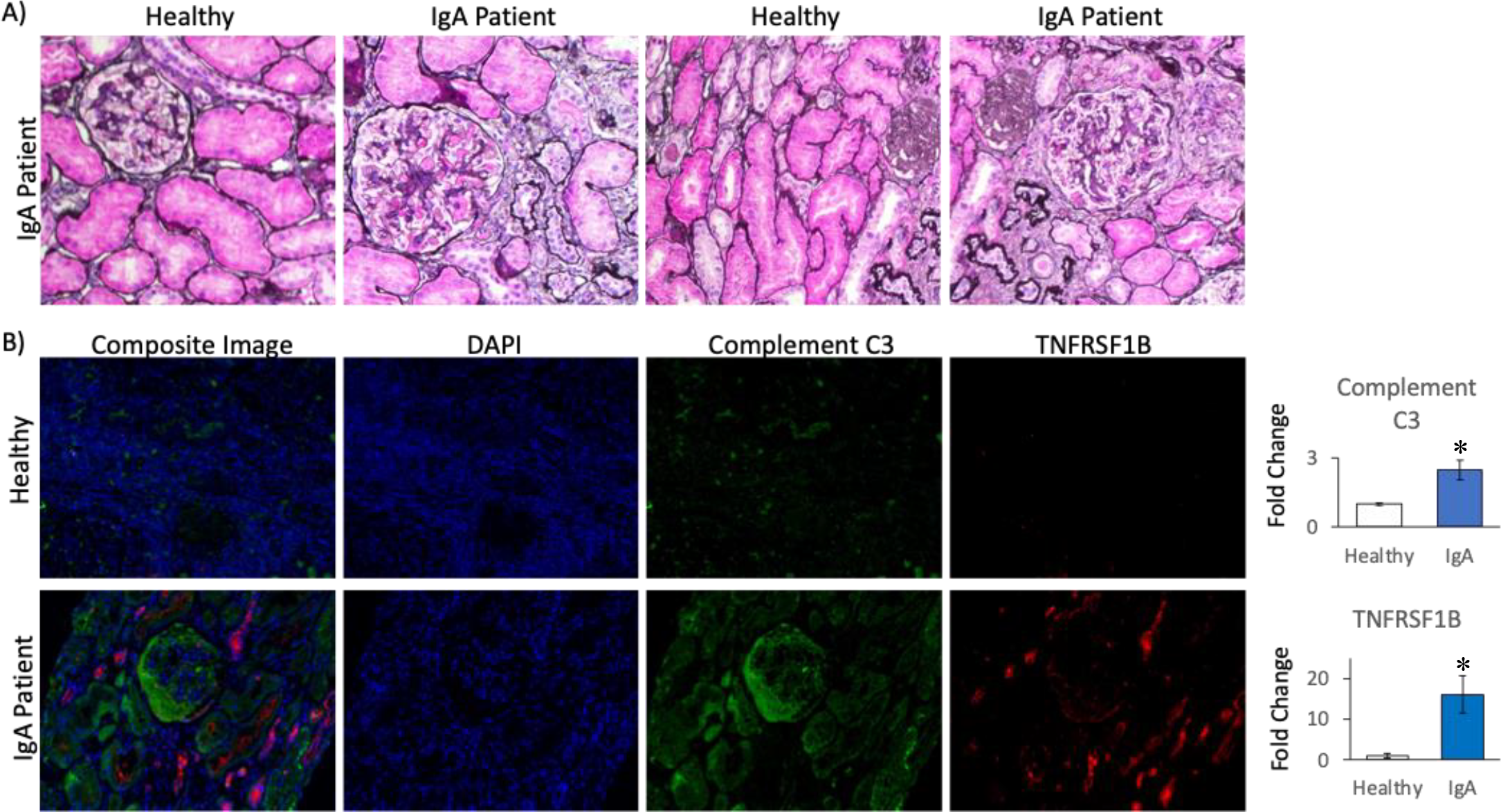
Kidney biopsy findings. (A) Light microscopy shows increased fibrosis in IgA nephropathy patients compared to healthy controls. (B) Representative immunofluorescence staining for Complement C3 and TNFSF1B in IgA nephropathy patients compared to healthy controls. Graph represents quantification of fluorescence intensity, * p<0.05).

### NanoString Identified mRNA Expressed Targets as Predictors of Fibrosis

Fibroblasts are responsible for the production and deposition of extracellular matrix components and may be stimulated by the persistent inflammation and immunological response, such as those during IgAN. The excessive production and accumulation of collagen and other matrix proteins lead to the formation of scar tissue, known as fibrosis. Therefore, we analyzed our IgAN patient samples for fibrosis formation and compared the severity of fibrosis with the Nanostring mRNA transcript expression of our targets. From our histological slides, we observed fibrosis formation in IgAN patients that resulted in structural changes within the kidney, including thickening of the glomerular basement membrane, obliteration of the glomerular capillaries, and an increase in the volume of the interstitial compartment (Fig. 1A). From our NanoString mRNA expression profiles, we observed a significant increase in *PTEN, CASPASE 3, TGM2, TGFB1, IL2*, and *TNFRSF1B* from IgAN patients with level 2 fibrosis compared to those with none (Table 5). In order to gain insights into such significantly upregulated mRNA from IgA fibrotic patients, we compiled them into the STRING platform (https://string-db.org/) to analyze any potential biological processes and interactions in the form of an interaction network. Set at a high confidence of 0.7, our identified “fibrotic” genes did not show any string correlation amongst themselves. However, when adding 5 more relevant nodes to the network, *IL2RB, SLC9A3R1, TP53, TGFBR2* and *XIAP*, we were able to establish a more complete network of associating genes, with a total of 11 nodes and 14 edges (Fig. 2). The protein-protein interaction networks (PPI) enrichment p-value of 0.0393 indicates that the nodes are not random and that the observed number of edges is significant. When studying the various descriptions in the “Biology Process” category, we identified that such genes are primarily involved in apoptotic pathways, regulation of apoptosis, wound healing, response to growth factors and regulation of lymphocyte activation, while the meaningful “Pathways” included Th17 cell differentiation, Cellular senescence, MAPK signaling pathway, and PI3K-Akt signaling pathway. Therefore, our study identified several upregulated mRNA transcripts that were upregulated during the development of fibrosis and may be considered as fibrotic markers within IgAN patients.

**Table 5.** Gene expression in biopsies of patients with stage 0 versus stage 2 fibrosis.

| Genes | Detected RNA transcripts |  |  |  | Ratio | P-value |
| --- | --- | --- | --- | --- | --- | --- |
|  | Stage 0 | SD | Stage 2 | SD |  |  |
| *PTEN* | 421.24 | 92.77 | 554.67 | 84.27 | -1.32 | 0.0084 |
| *CASPASE 3* | 78.76 | 18.00 | 109.64 | 28.43 | -1.39 | 0.0106 |
| *TGM2* | 191.84 | 121.35 | 353.69 | 165.28 | -1.82 | 0.0413 |
| *TGFB1* | 235.89 | 146.21 | 456.21 | 128.66 | -1.93 | 0.0087 |
| *IL2* | 1.29 | 0.54 | 2.64 | 2.41 | -2.05 | 0.0247 |
| *TNFRSF1B* | 75.83 | 52.50 | 158.20 | 44.83 | -2.09 | 0.0097 |

RNA, ribonucleic acid; SD, standard deviation.

**Figure 2.**
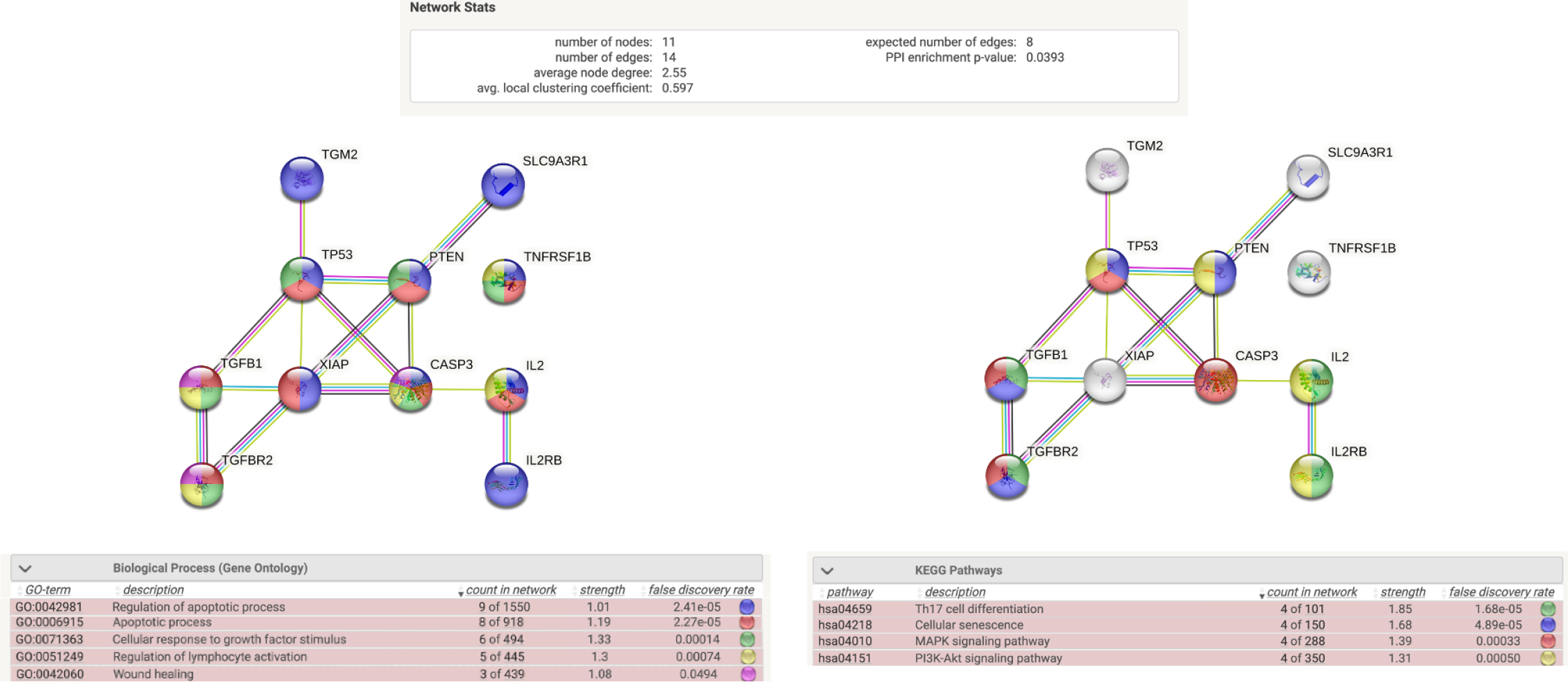
Mapping of associated fibrotic genes

## DISCUSSION

Our pilot study aimed to validate the use of the NanoString technology in the identification of potential inflammatory targets at different stages of IgAN disease progression. IgAN patients appears to have a distinct inflammatory profile than other inflammatory glomerulonephritis. We were able to identify *C3* and *TNFR2* as potential biomarker of early inflammation in patient with moderately increased risk of disease progression. When compared to healthy controls and using immunofluorescence technique, protein expression for those two markers was more pronounced, with distinct signal patterns. Those biomarkers might be implicated in the development of fibrosis when studying potential biological processes and interactions in the form of an interaction network.

Fibrosis is a common phenomenon of IgAN, which we confirmed within our study using Jones’ staining. In order to find an association of our significantly upregulated genes with the onset of fibrosis, we compared the expression of such genes between IgAN patients with grade 2 fibrosis to those without fibrosis. As a result, we identified 6 genes that were significantly associated with fibrosis: *PTEN, CASPASE 3, TGM2, TGFB1, IL2*, and *TNFRSF1B*. Interestingly, such genes were predicted to be associated with the biological functions of wound healing, apoptosis, cellular response to growth factors and regulation of immune systems. Fibrosis is a part of the normal wound healing process. When tissues are injured, a complex series of events takes place to repair the damage. Initially, there is an inflammatory response to remove debris and pathogens from the site of injury. Following this, various growth factors and cytokines are released, stimulating cell proliferation and migration to the wound site. Fibroblasts are activated and differentiate into myofibroblasts, which are the primary cells expressing excessive amounts of collagen and other extracellular matrix proteins. In fibrotic conditions, such as in chronic injuries or persistent inflammation, the wound healing process becomes dysregulated, leading to excessive fibrosis. Apoptosis, otherwise known as programmed cell death, can cause the loss of specific parenchymal cell types, including epithelial cells, in affected tissues. This cell loss contributes to the disruption of the normal tissue architecture, requiring tissue repair. Continuous apoptosis that requires chronic repair is a characteristic feature of fibrosis. Moreover, apoptotic cells release signals that can activate fibroblasts into myofibroblasts, which are the primary cells expressing large amounts of extracellular matrix components, leading to the formation of scar tissue. The immune system plays a crucial role in the development and resolution of fibrosis. In the initial stages of tissue injury or inflammation, immune cells, such as macrophages and lymphocytes, are activated and recruited to the site of injury. These immune cells release various cytokines and growth factors that can promote fibroblast activation and collagen deposition. Additionally, immune cells can directly interact with fibroblasts and modulate their function. In fibrotic diseases, the immune response may become dysregulated, leading to chronic inflammation and persistent fibrotic processes. Imbalances in immune cell populations and dysregulated immune signaling can contribute to the progression of fibrosis. Therefore, it is quite fitting that the predicted biological functions of our network of fibrotic IgAN genes are known to be important to fibrosis.

We also predicted various relevant pathways belonging to our IgA nephropathic genes that are significantly upregulated during fibrotic tissue formation. These include Th17 cell differentiation, cellular senescence, and the MAPK and PI3K-Akt signaling pathways. It has been shown that the abnormal humoral immunity observed in IgA nephropathy may be mediated by the differentiation of T helper 17 cells [9] and kidney senescence is well documented to occur after injury [10]. It has been observed that Akt activation is increased in experimental tubulointerstitial fibrosis [11, 12]. Akt activation serves as a crucial component in multiple signaling pathways associated with kidney damage. Consequently, there is a belief that Akt plays a significant role in renal fibrosis. The MAPK pathway plays a pivotal role in regulating both inflammatory and fibrotic responses within the kidney [13, 14]. Notably, one significant outcome of MAPK pathway activation is the induction of extracellular matrix protein synthesis, particularly collagen. The excessive deposition of collagen and other extracellular matrix components contributes to the progressive accumulation of fibrotic tissue in the kidney. Thus, the MAPK pathway serves as a key mediator in the development and progression of renal fibrosis by modulating various cellular and molecular events involved in fibrotic remodeling. These findings support the potential involvement of these pathways in driving the fibrotic process and provide valuable directions for further exploration.

Unfortunately, we failed to identify some biomarkers associated with a higher score at the IgA Nephropathy Prediction tool. Some biomarkers, known as *C1QA, ICAM1, TNF* and *TNFSF1B*, showed a distribution in function of the IgA prediction tool score range near a statistically significant level. Both *ICAM1* and *TNF* were close to be associated with a score range of over 20% at the IgAN prediction score. Presence of high levels of *ICAM-1* mRNA was associated in previous studies on IgAN with advanced histologic changes and a lower kidney function [15]. TNF-alpha is also suspected to play an important part in the pathogenesis of the IgA nephropathy. Previous study established a relation between high serum levels of TNF-a in patients with IgAN and MEST-C score, higher proteinuria and lower kidney function [16].

Our present study has some limitations. First, the small number of cases included reduced the power of our study and reduced the chances we had to find statistically significant differences in the distribution of the biomarkers analysis with the Nanotring technology for the clinically relevant endpoints we tried to focus on. However, the purpose of the study was not elaborate a prediction model using inflammatory biomarkers, but to validate Nanostring technology as a potential tool to delineate inflammatory processes in IgAN patients with ongoing kidney injury. Another limitation of our study is the limited number of candidate genes examined, mainly because of technical reasons. Nonetheless, we were able to focus on the main targets already described in the literature as involved in IgAN physiopathology.

In conclusion, our study validates the use of the NanoString technology as a promising tool for gene expression analysis in IgAN. By utilizing this innovative platform, we were able to simultaneously study multiple inflammatory biomarkers involved in IgAN and identify their activation patterns. We also validated the activation of genes that are known to contribute in kidney fibrosis, therefore shedding light on the underlying mechanisms of disease progression. Although these findings confirm the usefulness of gene expression profiles in providing useful tools for diagnosis and monitoring IgAN, further research is necessary to elucidate the clinical utility of gene expression analysis and its integration into predictive model s and treatment strategies for IgAN. Further exploration of gene expression profiles holds promise to a better comprehension of disease and the development of targeted therapies to mitigate kidney fibrosis in IgAN patients.

## Acknowledgments

The authors wish to thank Dr. Virginie Royal for providing the light microscopy biopsy images.

